# Enhancing cryopreservation of human iPSCs: Bottom-up vs Conventional freezing geometry

**DOI:** 10.1101/2024.10.30.621091

**Authors:** Fernando Teodoro, Soukaina El-Guendouz, Rafaela Neves, Andreia Duarte, Miguel A. Rodrigues, Eduardo P. Melo

## Abstract

Induced pluripotent stem cells (iPSCs) hold large potential on regenerative medicine due to their pluripotency and unlimited self-renewal capacity without the ethical issues of embryonic stem cells. To provide quality-controlled iPSCs for clinical therapies, it is essential to develop safe cryopreservation protocols for long-term storage, preferable amenable for scale-up and automation. We have compared the impact of two different freezing geometries (bottom-up and conventional radial freezing) on the viability and differentiation capability of human iPSCs. Our results demonstrate that the bottom-up freezing under optimized conditions significantly increases iPSCs viability, up to 9% for the cell membrane integrity and up to 21% for the cell metabolic state, compared to conventional freezing. The improvement achieved for bottom-up versus conventional freezing was maintained after scale-up from cryogenic vials to 30 mL bags, highlighting the method’s potential for clinical applications. These findings show that bottom-up freezing can offer a more controlled and scalable cryopreservation strategy for iPSCs, promoting their future use in regenerative medicine.

## Introduction

Clinical application of stem cells to treat human diseases has started in the 50s, with bone marrow-derived cells (Hunt, 2011). The haematopoietic stem cell source, whether from bone marrow, umbilical cord or, predominantly today from mobilised peripheral blood, is the most common source for cell-based therapies. More recently, a non-haematopoietic stem cell population, known as mesenchymal stem cells, has been subject to both laboratorial and clinical studies (Pittenger et al., 2019). Although, haematopoietic and non-haematopoietic adult stem cells show both multipotency and expansion potential, they do not form stable cell lines in culture, which is a limitation to breadth their clinical application, particularly in the field of regenerative medicine. On contrary, embryonic stem cells and induced pluripotent stem cells (iPSCs) can form stable cell lines with unlimited capacity for self-renewal and full pluripotent capacity to differentiate into cell types from all three germ layers (Hunt, 2011). This makes them of special significance in both regenerative medicine and toxicology. iPSC in particular, may broaden their utility without the ethical issues surrounding embryonic stem cells. The reprogramming of human somatic cells from patient-specific cells to an embryonic stem cell-like pluripotent state has the potential to be used in a variety of disease states without the need for immunosuppressants, as autologous transplantation of genetically identical cells overcomes the issue of immune rejection (Kiskinis and Eggan, 2010). Importantly, in case of a known genetic defect, gene therapy approaches could, in principle, restore normal cellular function. iPSCs constitute an unlimited supply of normal, differentiated cells with applications into toxicology and drug discovery (Csöbönyeiová et al., 2016; Rispoli et al, 2024), tissue engineering (Bastami et al., 2017) as well as gene and cellular therapy for a wide range of human diseases (Ilic and Ogilvie, 2022; Cecerska-Heryć et al., 2023).

To provide safe, quality-controlled iPSCs for clinical therapy and regenerative medicine, it is essential to develop robust cryopreservation protocols, as well as good manufacturing practices. Cell lines maintained in serial culture are susceptible to genetic variation and contamination, while the establishment of frozen banks of homogeneous populations can capture a single desired phenotype free of contamination. While vitrification is commonly used for long-term storage, involving high concentrations of cryoprotectants and storage in liquid nitrogen (-196°C), it has limitations such as high operational costs, safety concerns, and the toxicity of the cryoprotectants used (Hunt, 2011). The dependence on liquid nitrogen for cell storage increases operational expenses and raises issues related to impaired working efficiency and safety (Yuan et al., 2016). Also, cryopreservation of iPSCs by vitrification requires much higher concentrations of cryoprotectants than are usually tolerated by cells (Hunt, 2011). Another widely adopted conventional method for cryopreservation is radial freezing, typically used for short-term storage at -80°C. This method can be performed without controlled cooling rate, but more commonly, passive freezing devices are employed to achieve a rate of -1 °C/min, or active devices allow for adjustable freezing rates. The conventional methods rely on the use of cryoprotectants, typically dimethyl sulfoxide (DMSO), often at a 10% concentration. However, DMSO is toxic to cells and requires additional washing steps post-thaw to remove it, leading to significant cell loss and reduced viability. Although radial freezing at -80°C offers cost benefits and ease of use compared to vitrification, optimizing the freezing process to minimize DMSO toxicity and improve cell recovery after thawing remains a challenge. Developing innovative cryopreservation techniques and media formulations is crucial to enhance iPSC viability without compromising pluripotency for clinical applications.

In the cryopreservation process, the direction of ice growth is crucial for ensuring quality and reproducibility. Directional freezing, which allows for a more precise control over ice growth and morphology, has demonstrated positive outcomes in various studies (Bahari et al., 2018; M. A. Rodrigues et al., 2013; Si et al., 2010). For instance, Bahari et al. (2018) introduced a controlled slow cooling technique combining initial directional freezing with gradual cooling to -80°C. Optimal post-thaw viability of Caco-2 cells at approximately 70%, compared to only 15% with non-directional freezing was achieved. Bottom-up (unidirectional) freezing offers a novel approach for cryopreservation, allowing more precise control over ice growth, thereby potentially improving cell viability. In this work we have studied the impact of bottom-up slow-cooling to -80°C on the viability and differentiation capability of human iPSCs into neurons. We tested different variables using the bottom-up freezing geometry, namely DMSO concentration, ice nucleation time, and cooling rate on the viability and differentiation capability of iPSCs, always in parallel with conventional freezing at -80°C. We have also scaled-up the freezing volume from 1 mL vials to 30 mL bags using bottom-up freezing in comparisson with conventional freezing.

## Results and Discussion

We have measured the viability and differentiation of human iPSC into mature neurons to compare freezing cryogenic vials at -80°C with two different geometries, conventional and bottom-up (unidirectional). Frozen storage of iPSCs at -80°C is possible for at least one year without significantly compromising cell viability and pluripotency (Yuan et al., 2016). The cell line used was a stable clonal transgenic line of human iPSC harbouring the doxycycline-inducible transcription factor neurogenin-2 at a safe harbour loci (Wang et al., 2017). Transcription factor overexpression is a new approach to neuronal differentiation from iPSCs that circumvent many challenges associated with small molecule-based differentiation protocols, such as low efficiency, different rates of differentiation for individual cells and, different iPSC clones responding differently to the same small molecule (Fernandopulle et al., 2018). The iPSCs used can be differentiated with near 100% efficiency and purity, and in a simple two-step protocol, into mature glutamatergic cortical neurons in the presence of doxycycline. For conventional freezing, the CoolCell freezing container (from Corning) reported to provide a freezing rate of -1°C/min when placed at -80°C was used. For bottom-up freezing with controlled ice-nucleation, the CELL equipment (from SmartFreez) opens new possibilities to optimize freezing parameters. Firstly, a layer of nucleation ice at the bottom of the cryogenic container can be formed for a short time at -80°C and then the bulk of cell suspension can be frozen (from bottom to top) from -5°C to -80°C at a variable freezing rate. This bottom-up freezing strategy allows a more precise control of the ice growth velocity. After two days frozen at -80°C, the cells were thaw at 37°C and viability was measured through membrane integrity using trypan blue. iPSCs viability was also evaluated through a metabolic assay, which correlates with the reducing power of living cells, applied on adherent cells, 24 h after thawing (see standard curve in figure S1). Optimization of freezing parameters based on cell membrane integrity only might be inadequate as the ability of iPSCs to attach to culture substrates (plating efficiency) does not correlate with membrane integrity for a wide range of freezing parameters (Li et al., 2018; Yuan et al., 2016).

### iPSC viability after freezing with different percentages of dimethyl sulfoxide (DMSO)

DMSO is known to be toxic to cells (Galvao et al., 2014). In patients receiving cryopreserved stem cells, DMSO can lead to adverse reactions. Therefore, reducing DMSO concentrations has been reported to alleviate these adverse reactions, improving the safety of stem cell therapies (Berz et al., 2007). Different percentages of DMSO in the freezing media were tested under conventional and bottom-up freezing. Conventional freezing was performed at a rate of -1 °C/min to -80 °C using the CoolCell freezing container. The same conditions were applied using the bottom-up freezing CELL equipment, but firstly a layer of nucleation ice was created at the bottom of the cryogenic tube (through the contact with a contact fluid at -80 °C for 90 s, see material and methods) before freezing the bulk cell suspension at a rate of -1 °C/min. The cell viability measured through membrane integrity using trypan blue is clearly higher than the one measured through the metabolic assay (Figure 1). This might result from dead cells that keep membrane integrity (do not internalise trypan blue but as dead cells do not adhere and not respond to the metabolic assay). Plating efficiency was reported to be lower compared to membrane integrity (Yuan et al., 2016). It may also result from live cells that adhere to the culture substrate but are unable to respond to the metabolic assay if the onset of full metabolic activity for some live cells takes more than 24 h after thawing, when the metabolic assay was applied to. Apoptosis may also cause some early cell death after cell plating, decreasing the number of cells that respond to the metabolic assay, as documented for human embryonic stem cells (Heng et al., 2006; Xu et al., 2010). As expected, decreasing DMSO to 1% compromises significantly cell viability for both freezing geometries and both viability methods. Cell viability is similar for conventional and bottom-up freezing, except for membrane integrity at 1% DMSO where bottom-up freezing leads to a 4-fold increase on iPSCs viability (from 5 to 20%) (Figure 1A). This result indicates that bottom-up freezing is mechanically less aggressive. While in 10% DMSO the general porosity of the frozen matrix is high, decreasing DMSO concentration compacts the structure with higher ice content, therefore intensifying the mechanical stresses of the non-crystalizing phase containing the cells. Bottom-up unidirectional freezing favours the alignment of ice dendrites and prevents the enclosure of unfrozen liquid, which can raise internal pressure as water expands with crystallization (Duarte et al., 2020). Notwithstanding the larger value for the membrane integrity, the cell metabolic state was equally compromised, showing no significant differences between both freezing geometries (Figure 1B).

**Figure 1.**
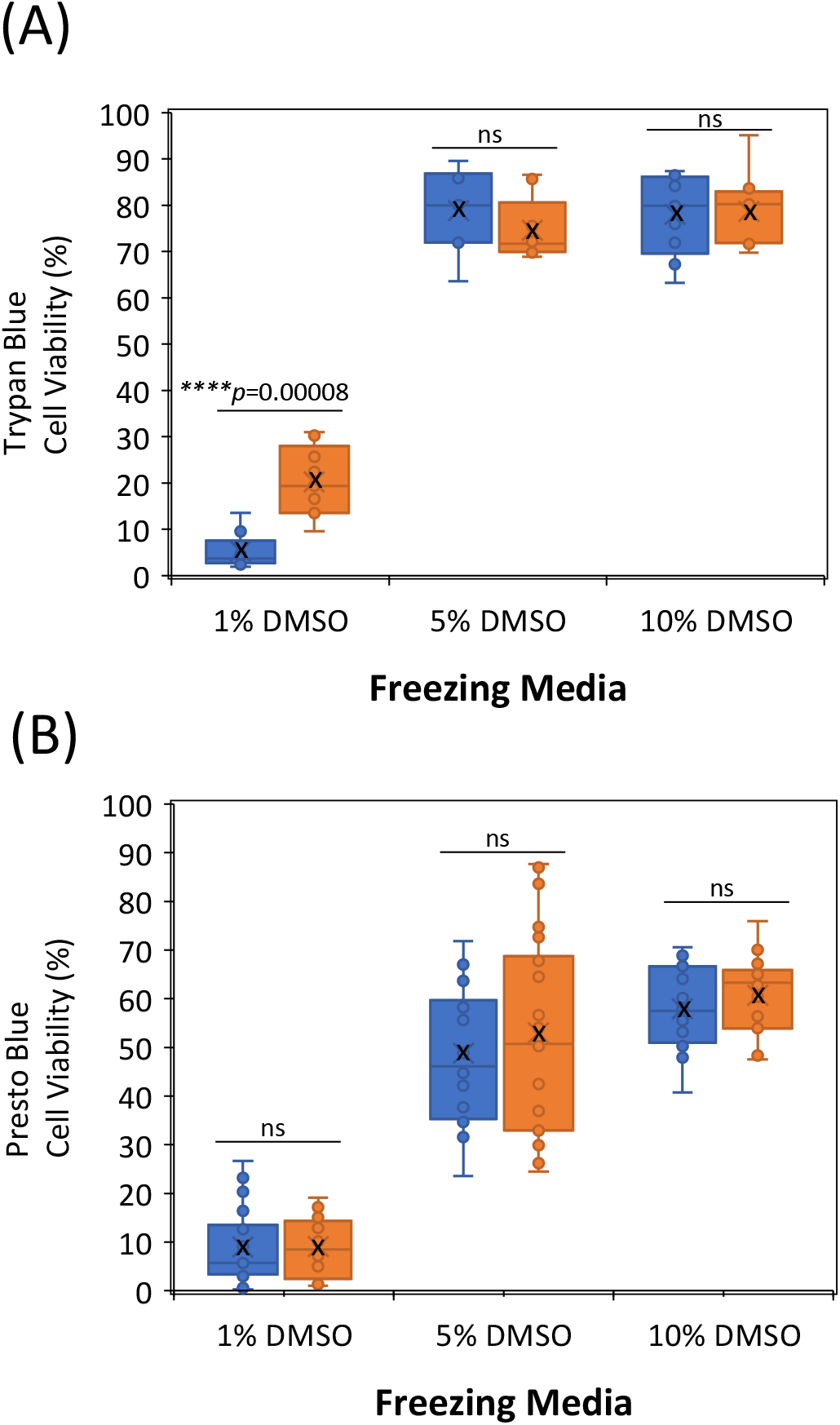
Viability of human iPSC after conventional (blue boxes) or bottom-up freezing (orange boxes) with different percentages of DMSO measured through membrane integrity (A) and through the cell reducing power (B). Each data point is an independent biological replicate (independent freezing tube) or an independent experimental replicate (independent cell plating well in the case of cell reducing assay). The average cell viability for each condition was highlighted by a black cross. \*\*\*\**p*<0.0001; ns: not significant for *p*>0.05; one way ANOVA statistical analysis

### iPSC differentiation into neurons

Differentiation of iPSCs into mature neurons last for 8 days in the presence of doxycycline (Figure 2). Neuronal morphogenesis with the onset formation and outgrowth of axons and dendrites is visible at day 5 and extends to day 8 for fully differentiation. To evaluate differentiation pace and degree, we have measured the length of terminal branches and number of branch nodes per cell from day 5 to day 8, using the simple neurite tracer FIJI plugin (Figure 3). Differentiation occurs at similar pace and cells reach fully differentiation after conventional and bottom-up freezing for the three DMSO concentrations tested. This shows that the freezing geometry and amount of DMSO do not affect the capability of live iPSCs to differentiate into neurons.

**Figure 2.**
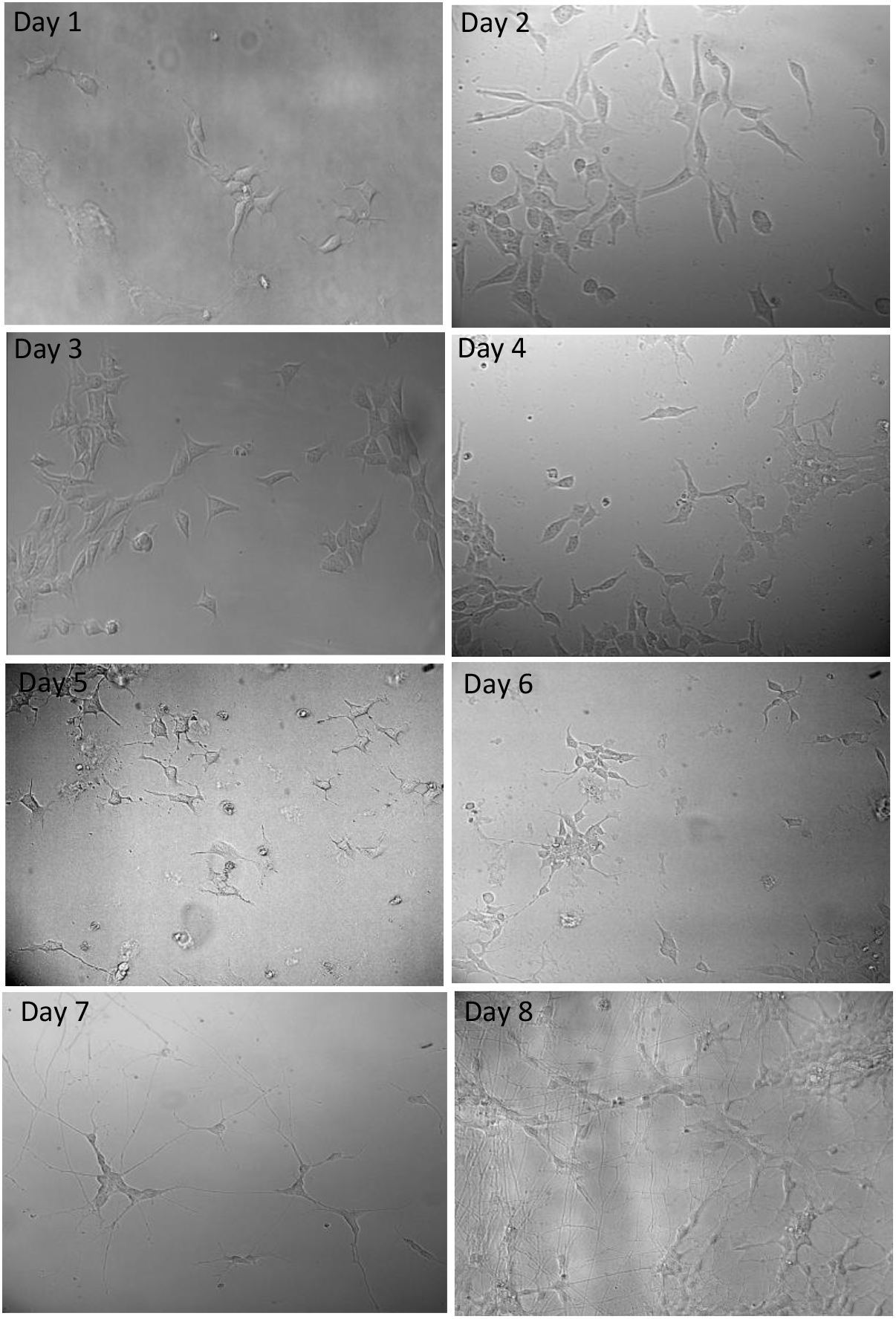
Images from the time course of iPSC differentiation into mature neurons.

**Figure 3.**
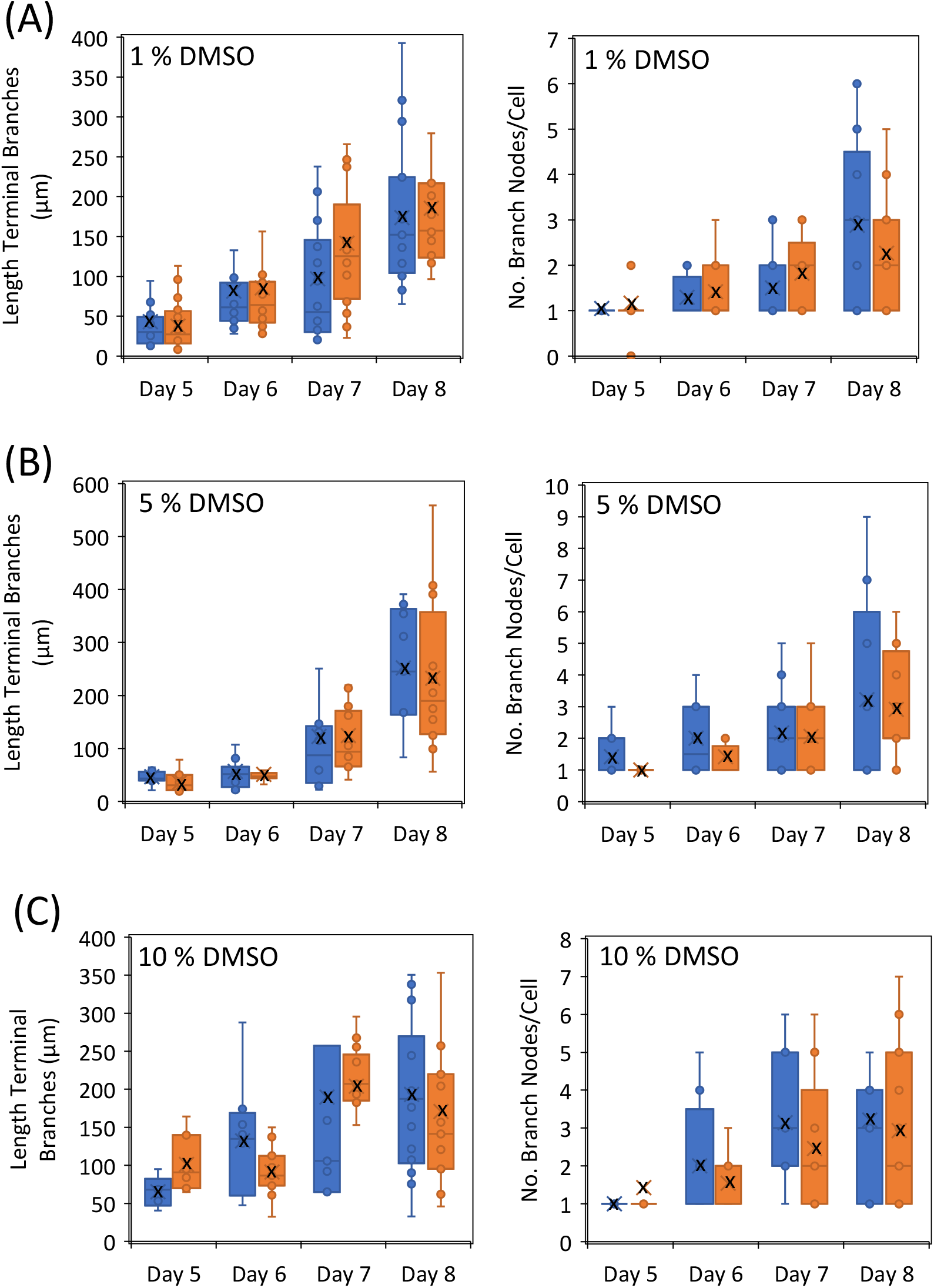
Evaluation of iPSC differentiation into neurons after conventional (blue boxes) or bottom-up freezing (orange boxes) with different percentages of DMSO (1 % in (A); 5 % in (B) and 10 % in (C)). The Simple Neurite Tracer plugin in FIJI package was used to quantify the lenght of terminal branches (left plots) and the number of branch nodes per cell (right plots). The average value for each condition was highlighted by a black cross.

### iPSC viability after bottom-up freezing with variable ice nucleation time

During slow cooling of a cell suspension, ice nucleation is initiated in the extracellular compartment, with cells present within the solute channels that form between the growing ice crystals. For bottom-up freezing, a thin layer of ice is formed at the bottom of the cryogenic tube to permit a controlled growth of the ice matrix with ice dendrites growing aligned from the bottom (ice layer) to the top of the cryovial. This ice layer should be as thin as possible (preferably less than 10% of the total volume) because it forms in an uncontrolled manner and the cells entrapped in this initial layer can experience mechanical stresses. To study this effect, the nucleation ice layer was formed at -80 °C for different nucleation times (60, 90 and 120 s), and then the bulk of cell suspension was frozen from -5 to -80 °C at a freezing rate of -1 °C/min (Figure 4). A concentration of 5% DMSO was selected to carry out this study to use a lower amount of DMSO without significantly compromising cell viability, as observed for 1% DMSO. Cell viability measured through membrane integrity is larger than measured through the metabolic assay as reported before in Figure 1. Variable nucleation times do not affect significantly viability measured through membrane integrity and values are similar for the two freezing geometries (Figure 4A). However, viability measured through the reducing power of living cells showed a clear dependence of the nucleation time (Figure 4B). Average cell viability was 46% for 60 s nucleation time, increased to 53% for 90 s, but is highly compromised for 120 s with only 19% average cell viability. This reduction in cell viability for 120 s may be associated with the larger amount of ice formed uncontrolled and too rapidly during the nucleation phase, potentially reducing the amount of bulk suspension frozen under controlled bottom-up freezing. For 90 s nucleation time, there was a statistically significant net increase on cell viability from 41% with conventional freezing to 53% using bottom-up topology. Optimization of the nucleation time to 90 s permitted a 1.3-fold increase on average cell viability measured through the metabolic assay compared to conventional freezing. Worth notice, the larger scatter of cell viability values measured through the metabolic assay under conventional freezing (Figure 4B), reflecting probably a more uncontrolled freezing and stochastic expansion of the ice matrix in the absence of a seed of nucleating ice. The pace and degree of iPSC differentiation into neurons was not affected by using different nucleation times (Figure S2).

**Figure 4.**
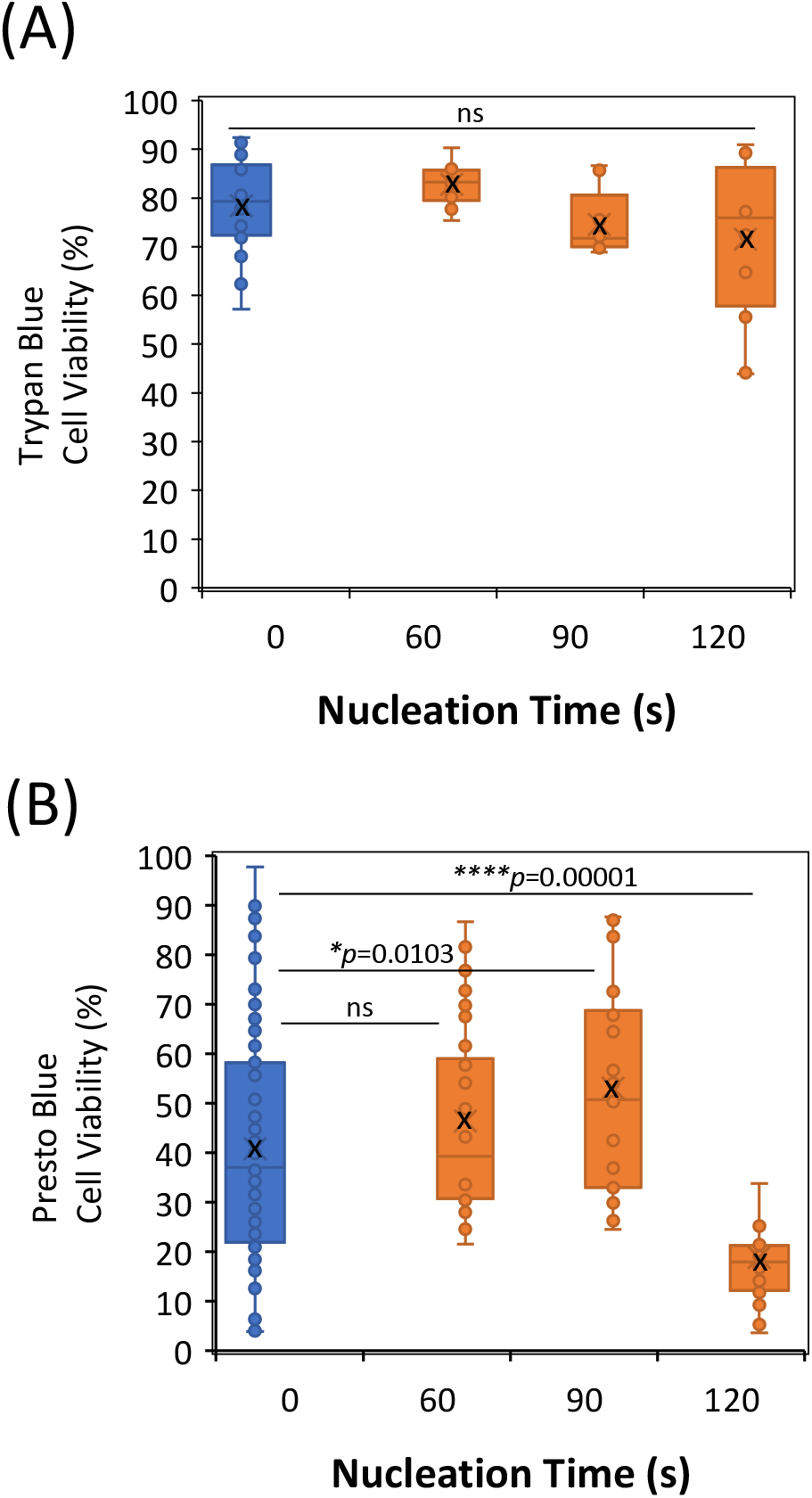
Viability of human iPSC after conventional (blue boxes) or bottom-up freezing (orange boxes) with 5 % DMSO, measured through membrane integrity (A) and through the cell reducing power (B). Different nucleation times were applied for bottom-up freezing, always in parallel with an independent conventional freezing experiment. Each data point is an independent biological replicate (independent freezing tube) or an independent experimental replicate (independent cell plating well in the case of cell reducing assay). The average cell viability for each condition was highlighted by a black cross. \**p*<0.05 \*\*\*\**p*<0.0001; ns: not significant for *p*>0.05; one way ANOVA statistical analysis

### iPSC viability after bottom-up freezing with different cooling rates

Another freezing parameter known to affect cell viability is the cooling rate (Li et al., 2018), easily controlled for bottom-up freezing. With 5% DMSO and 90 s nucleation as the best conditions revealed in Figure 4, we have tested different cooling rates under bottom-up freezing in comparison with conventional freezing at -1 °C/min provided by the CoolCell container. Again, cell viability measured through membrane integrity is larger than measured by the metabolic assay (Figure 5). Increasing the cooling rate from -1 °C/min to -2 °C/min under bottom-up freezing increases the average cell viability from 75% to 87% regarding membrane integrity, though not statistically significant (Figure 5A). The metabolic assay revealed a statistically significant increase from 53% to 62% for the same increase in the cooling rate (Figure 5B). Further increase of the cooling rate to -3 °C/min compromised the average cell viability slightly. The pace and degree of iPSC differentiation into neurons was not affected by different cooling rates (Figure S3).

**Figure 5.**
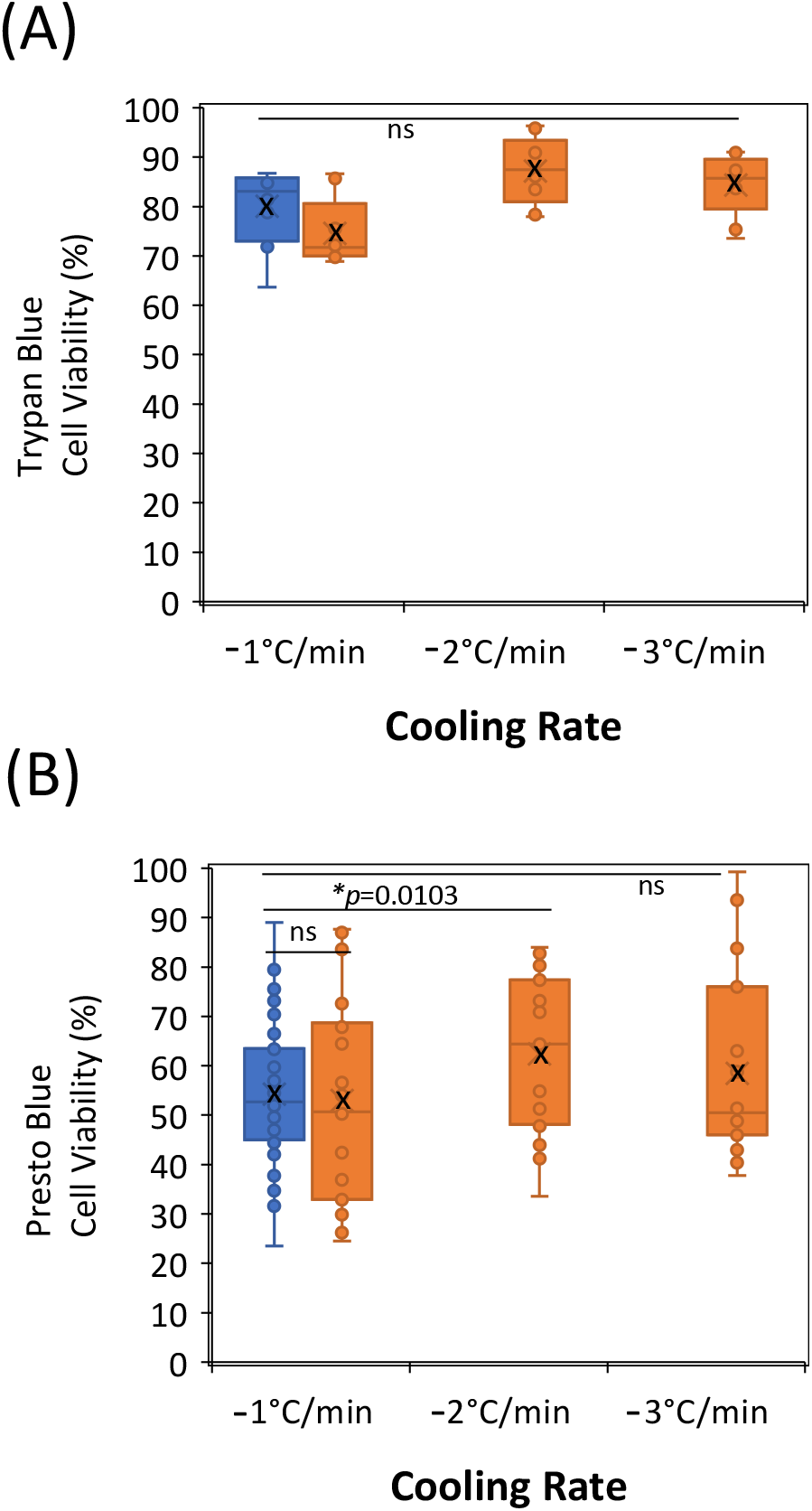
Viability of human iPSC after conventional (blue boxes) or bottom-up freezing (orange boxes) with 5 % DMSO and 90 s nucleation time, measured through membrane integrity (A) and through the cell reducing power (B). Different colling rates were applied for bottom-up freezing, always in parallel with an independent conventional freezing experiment at -1°C/min provided by the CoolCell freezing container. Each data point is an independent biological replicate (independent freezing tube) or an independent experimental replicate (independent cell plating well in the case of the cell reducing assay). The average cell viability for each condition was highlighted by a black cross.

### Scale-up of iPSC freezing

Cryopreservation methodologies amenable to scale-up are needed (Hunt, 2011). The data shown in previous figures was obtained using 2 mL cryogenic tubes with 1 mL cell suspension (see material and methods). To test the scale-up of iPSC cryopreservation we have used 30 mL bags with 5% DMSO in the freezing media (Figure 6). For conventional freezing the bag was placed at – 80 °C, and for bottom-up freezing, a layer of nucleation ice was formed for 90 s at – 80 °C and then the cell suspension was cooled from -5 °C to – 80 °C at -2 °C/min cooling rate, chosen from the previous results. Scale-up compromises iPSC viability, with around 20% decrease on viability measured through membrane integrity and a 51% decrease measured through the cell reducing power, for both conventional and bottom-up freezing (compare Figures 5 and 6). Comparing viability for the two freezing geometries after scale-up revealed better values for bottom-up freezing, with 69% compared to 60% for membrane integrity and 31% compared to 23% for the cell reducing power.

**Figure 6.**
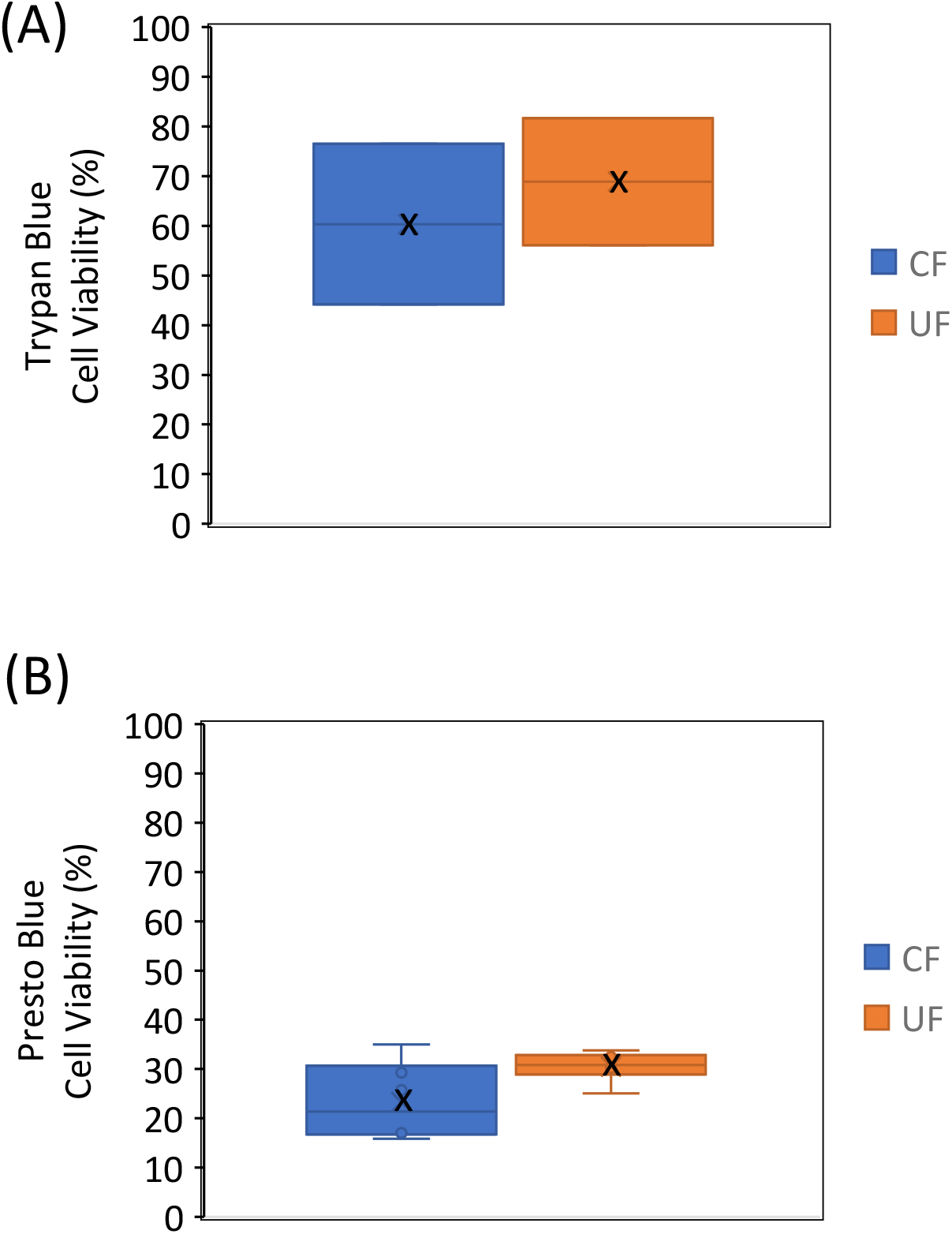
Scale-up for freezing iPSC in 30 mL bags. Viability of human iPSC after conventional (CF, blue boxes) or bottom-up freezing (UF, orange boxes) with 5 % DMSO, measured through membrane integrity (A) and through the cell reducing power (B). Conditions for bottom-up freezing were 90 s nucleation time and -2 °C/min cooling rate. The average cell viability for each condition was highlighted by a black cross.

## Conclusions

Cryopreservation of iPSCs as an undifferentiated cell suspension using simple procedures amenable for automation, asepsis and scale-up, might be of major importance for stem cells usage in different applications, particularly on clinical therapy. Bottom-up freezing offers new possibilities to optimize freezing parameters. Firstly, a layer of nucleating ice can be formed at the bottom of the freezing container to achieve a more controlled growth of the ice matrix. Secondly, heat-transfer is less stochastic using a bottom-up geometry, allowing a more effective control of the cooling rate and of the ice structure. The bottom-up alignment of ice dendrites resulted generally in less membrane rupture, due to the attenuation of mechanical stresses, which was more significant under 1% DMSO. Bottom-up freezing of iPSCs compared to conventional freezing, both at – 80°C, resulted on the improvement of average cell viability, particularly when measured through the cell metabolic state. Under optimized conditions, i.e., 90 s nucleation time and -2°C/min cooling rate, bottom-up freezing using 2 mL cryovials has resulted in 87% and 62% average cell viability for membrane integrity and cell reducing power, respectively. This is an increase on iPSCs viability of 7-9% based on membrane integrity and 8-21% based on the cell reducing power, compared to the conventional freezing. Moreover, the advantage of bottom-up versus conventional freezing was maintained after scale-up to 30 mL bags, with a 9% increase on viability measured through membrane integrity and an 8% increase through the cell reducing power. In sum, the results shows that the controlled bottom-up freezing could be a good strategy to improve the cryopreservation methods for cell therapies, allowing to reduce the amount of DMSO typically used, while maintaining a high post-thaw cell viability.

## Material and Methods

### iPSC culture

Vitronectin (Gibco, USA) was thawed on ice and diluted in Phosphate-buffered saline (PBS) at a 1:150 ratio to coat 70 mm cell culture dishes (Avantor, China) and 96-well plates (Thermo Scientific, Denmark). The coated plates were then incubated overnight at room temperature and the leftover vitronectin solution was removed before use. iPSCs were thawed in a 37°C water bath and immediately transferred to room temperature Essential 8 (E8) medium (Gibco, USA) to dilute the freezing media. The cells were then centrifuged at 1000 rpm for 3 minutes, and the supernatant was discarded. The cells were resuspended in fresh E8 medium supplemented with 10 μM ROCK inhibitor (Y-27632, BD Biosciences, USA) and seeded onto vitronectin-coated cell culture dishes. The presence of ROCK inhibitor for Rho kinases has shown to reduce markedly dissociation-induced apoptosis in stem cells (Watanabe et al., 2007). After 24 hours, the E8 medium containing the ROCK inhibitor was changed to E8 Flex medium (Gibco, USA) and the media was changed every day until passaging was required. Upon reaching approximately 70%-80% confluency, cells were washed with PBS and detached using 1 mL of Accutase (Invitrogen, Canada) for 3-5 minutes at 37°C and then quenched with E8 medium. The cells were then centrifuged at 1000 rpm for 3 minutes and resuspended in fresh E8 medium supplemented with 10 μM ROCK inhibitor before being seeded onto vitronectin-coated cell culture dishes or 96-well plate.

### iPSCs freezing

For cryopreservation of iPSCs stocks, cells were detached using Accutase, resuspended in the cryopreservation solution consisting of 40% E8 medium, 50% KnockOut Serum Replacement (KSR, Gibco, USA), 10% dimethyl sulfoxide (DMSO, Sigma-Aldrich, USA) and 10 μM ROCK Inhibitor, and then transferred to 2 mL cryovials (VWR, Canada). The cryovials were then placed in a CoolCell (Corning, USA) container and frozen at -80°C for 24 hours before being transferred to -150°C for long-term storage.

To assess the effect of the freezing media and freezing parameters on the viability of iPSCs, the cells were detached with Accutase, centrifuged, resuspended in fresh E8 medium, and a small aliquot was taken to stain with trypan blue (Merk, USA) and cell counting using a Coverslip-Microscope Slide (KOVA Plastics, USA). The bulk of cell suspension in E8 medium was then diluted in the freezing media to a cell density of 1.5-2×10^6^ cells/mL. To assess the effect of different percentages of DMSO, the composition of the freezing media was E8 medium at a percentage of (50% - % DMSO), 50% KnockOut Serum Replacement and 10 μM ROCK Inhibitor. The cell suspension in the freezing media was split into two groups (each composed by three cryovials), one for conventional freezing and the other for bottom-up freezing in the CELL controlled rate freezer equipment from SmartFreez, Portugal.

For conventional freezing, the cryovials were placed in the CoolCell container (reported to impose a cooling rate of -1 °C/min), frozen at -80 °C for 24 hours and then maintained at -80 °C for another 48 hours. For bottom-up freezing, the cryovials were placed in the pre-chilled holder and then in the CELL equipment. During the nucleation step, a thin layer of ice was formed at the bottom of the cryovials by touching down the vials into a contact fluid (ethanol) at -80°C for the specified time gap. This step is crucial to ensure the controlled bottom-up ice growth. The cryovials were then lifted up from the contact fluid to achieve the starting temperature of the freezing program, and then cooled from -5°C to -80°C at a specified cooling rate.

After freezing, the cryovials were quickly transferred to a -80°C freezer, where they remained for 72 hours. After this period, the cells from both groups were thawed in a water bath at 37°C and resuspended in E8 media containing the ROCK inhibitor. A small aliquot from each cryovial was taken for posterior cell counting using trypan blue staining. The remainder of the cell suspension of each cryovial was plated both on a vitronectin-coated 96-well plate and on a geltrex-coated coverslip at desired density. The vitronectin-coated 96-well plate was incubated at 37°C, and after 24h the E8 media was removed and 90 μL of E8 Flex media and 10 μL of PrestoBlue HS reagent (Invitrogen, USA) were added. The cells were incubated at 37°C for 4h before the absorbance at 570 nm and 600 nm were measured using a microplate reader (BioTek Synergy Neo2). The cells seeded in the geltrex-coated coverslip were used to study the process of differentiation into neurons.

### iPSCs differentiation into neurons

*Geltrex Coating:* Geltrex (Gibco, USA) was thawed in ice and diluted in DMEM/F12 (Gibco, Netherlands) at a ratio of 1:100. The μ-Slide 8-well coverslips (Ibidi, Germany) were coated with 200 μL per well of the diluted Geltrex solution and incubated at 37°C overnight. Prior to cell seeding, the leftover liquid was removed.

*Differentiation – Day 1 and 2:* A day after the cells have been seeded, the E8 media was removed, and replaced by iN1 media medium consisting of DMEM/F12 supplemented with N2 supplement (100x, Gibco, USA), L-Glutamine (100x, Gibco, UK), non-essential amino acids (100x, Biowhitaker, USA), 50 μM 2-Mercaptoethanol (Gibco, UK) and 1 μg/mL doxycycline (Thermo Fisher Scientific, USA). The medium was replaced by fresh iN1 every day until the end of the 2^nd^ day.

*Differentiation – Day 3 through 8:* At the start of the 3^rd^ day, the iN1 media was removed and iN2 media consisting of Neurobasal media (Gibco, UK) supplemented with L-Glutamine, B-27 supplement (50X, Gibco, USA), 10 ng/mL NT-3 (BRAND), 50 μM 2-Mercaptoethanol, 10 ng/mL BDNF (Thermo Fisher Scientific, USA) and 1 μg/mL of doxycycline was added. Full media change was performed every day until the 6^th^ day. No media change on the 7^th^ day and half of the media was replaced on the 8^th^ day.

### Scale-up of iPSC freezing

To assess the performance of a scale-up setting, 30 mL CryoMACS Freezing Bags 50 (Miltenyi Biotec, Germany) were used instead of 2 mL cryovials. The conditions upon bottom-up freezing were nucleation time 90 s, cooling rate -2°C/min and freezing media with 5% DMSO. For conventional freezing, the 30 mL bag was placed directly into the - 80°C freezer.

### Image acquisition and analysis

Differentiation of iPSCs into mature neurons was imaged on an Olympus IX71 microscope under bright field. To quantify the length of terminal branches and the number of branch nodes per cell during differentiation, the Fiji software (version ImageJ 1.53Q) and the Simple Neurite Tracer (SNT) plugin (version 4.0.13) were used for image analysis.

## Acknowledgements

This study received Portuguese national funds from FCT - Foundation for Science and Technology through project UIDB/04326/2020 (DOI:10.54499/UIDB/04326/2020), UIDP/04326/2020 (DOI:10.54499/UIDP/04326/2020) and LA/P/0101/2020 (DOI:10.54499/LA/P/0101/2020), from the operational programmes CRESC Algarve 2020 and COMPETE 2020 through projects EMBRC.PT ALG-01-0145-FEDER-022121 and, from PT2020 through project SMARTCELL (LISBOA-01-0247-FEDER-047169).

