## supplementary figures for "Enhancing cryopreservation of human iPSCs: Bottom-up vs Conventional freezing geometry"

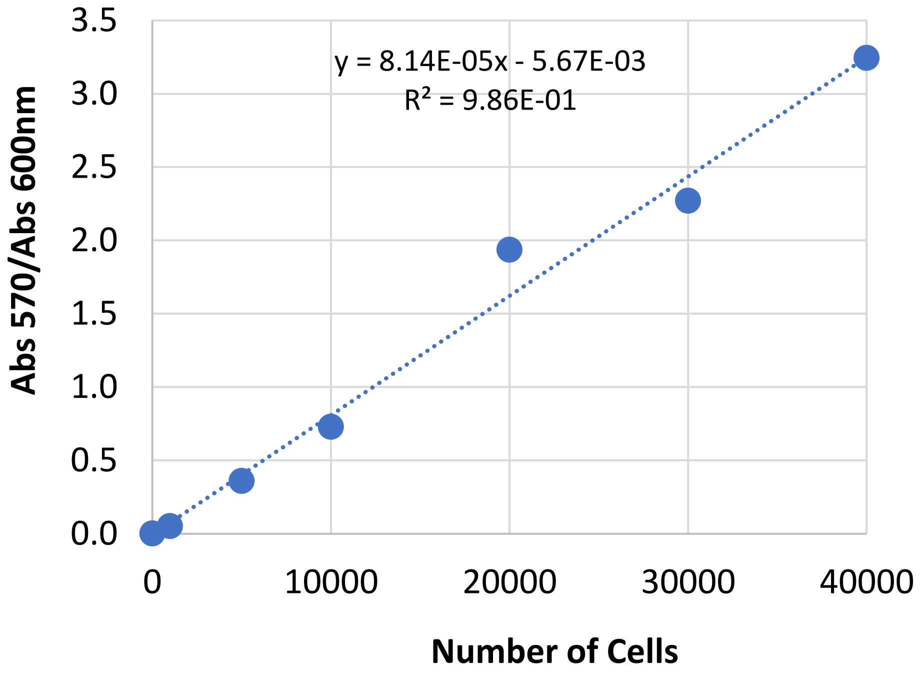


Figure S1. Standard curve for the response of Presto Blue (which correlates with the reducing power of living cells) versus number of adherent human iPSC.

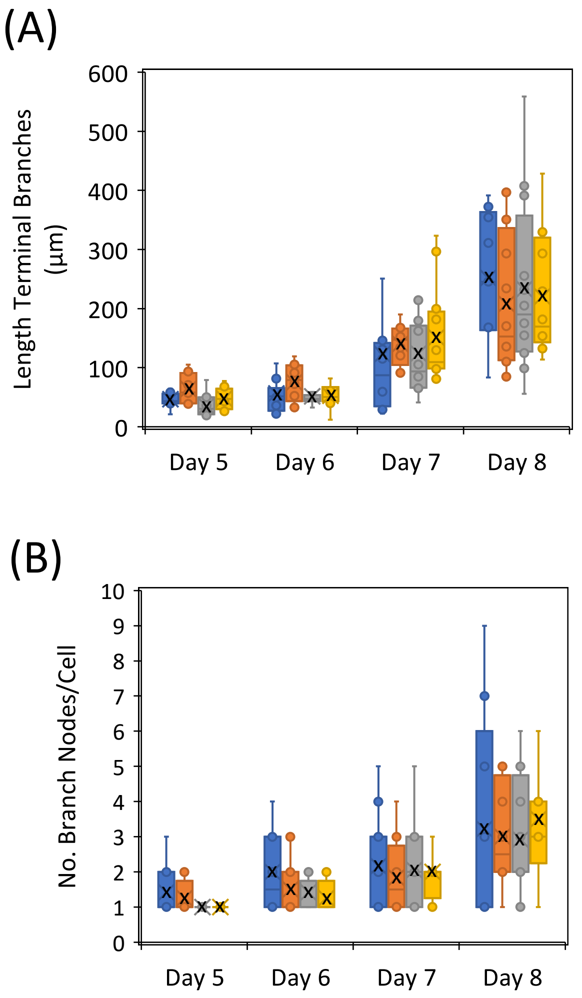


Figure S2. Evaluation of iPSC differentiation into neurons after conventional (blue boxes) or bottom-up freezing for different nucleation times (orange 60 s; gray 90 s; yellow 120 s). The Simple Neurite Tracer plugin on FIJI package was used to quantify the lenght of terminal branches (A) and the number of branch nodes per cell (B). The average value for each condition was highlighted by a black cross.

(A)

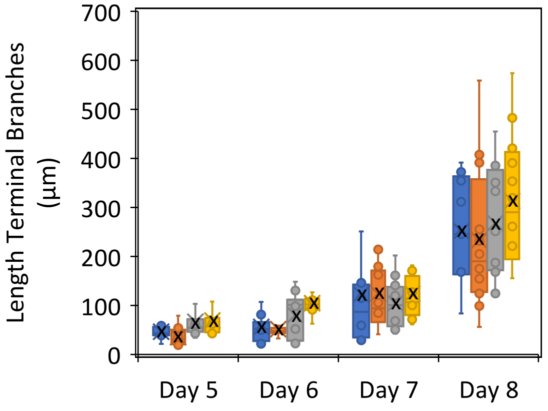


(B)

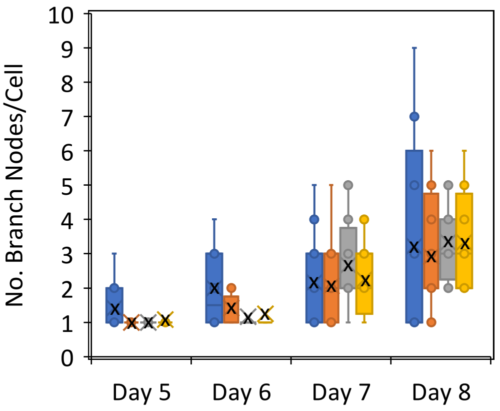


Figure S3. Evaluation of iPSC differentiation into neurons after conventional (blue boxes; -1 °C/min) or bottom freezing for different cooling rates (orange -1 °C/min; gray -2 °C/min; yellow -3 °C/min). The Simple Neurite Tracer plugin on FIJI package was used to quantify the lenght of terminal branches (A) and the number of branch nodes per cell (B). The average value for each condition was highlighted by a black cross.
